# Evaluation of Postharvest Senescence in Broccoli Via Hyperspectral Imaging

**DOI:** 10.1101/2020.12.16.423030

**Authors:** Xiaolei Guo, Yogesh Ahlawat, Alina Zare, Tie Liu

## Abstract

Fresh fruits and vegetables are invaluable for human health, but their quality deteriorates before reaching consumers due to ongoing biochemical processes and compositional changes. The current lack of any objective indices for defining “*freshness*” of fruits or vegetables limits our capacity to control product quality leading to food loss and waste. It has been hypothesized that certain proteins and compounds such as glucosinolates can be used as an indicator to monitor the freshness of vegetables and fruits. However, it is challenging to “visualize” the proteins and bioactive compounds during the senescence processes. In this work, we propose machine learning hyperspectral image analysis approaches for estimating glucosinolates levels to detect postharvest senescence in broccoli. Therefore, we set out the research to quantify glucosinolates as “freshness-indicators” which aid in the development of an innovative and accessible tool to precisely estimate the freshness of produce. Such a tool would allow for significant advancement in postharvest logistics and supporting the availability for high-quality and nutritious fresh produce.

## Introduction

Broccoli (*Brassica oleracea* L. var. *italica*) is a nutritious vegetable that is well enriched in anti-cancerous chemical compounds like glucosinolates [1]. Broccoli is highly perishable and senesces quickly after harvest. Broccoli is usually harvested at a developing stage of inflorescence therefore causes stress-induced senescence. The stress-induced senescence leads to faster chlorosis and increases in proteases resulting in the dismantling of chloroplast number and component [2]. Thus, during the postharvest storage and transportation, broccoli florets starts to turn yellow accompanied with a decrease in nutritional quality [1].

Senescence is a developmental process that can be tracked by monitoring physiological and biochemical changes in transcripts, proteins, and metabolites in broccoli. One particularly interesting class of phytochemicals are glucosinolates, given their importance not only for plant protection, but also for their dietary significance as chemo-preventative compounds that are found in edible plants, (i.e., cruciferous crops). However, the actual quality, storability and overall “freshness” of broccoli postharvest is quite uncertain unless the changes are visible to the human eye. Detecting any deteriorative physiological signs and symptoms before any irreversible damage occurs could allow for the development of freshness indicators, which can be used to identify the best postharvest handling, process and storage practices [3]. Therefore, sensitive indicators of potential deterioration are essential for extending optimal postharvest quality.

Prior work to detect symptoms of degradation in quality include using color measurements at an early postharvest stages [4]. Chlorophyll fluorescence and RGB (red, green, blue) color imagery analysis were used to measure the pigment change in broccoli using florescence and inverse red channel for color measurements [5]. However, objectively measuring the progressive loss of freshness after harvest has been a heretofore intractable problem in postharvest handling of fresh produce. Until now, determining freshness has been mostly based on external visual criteria like wilting, shriveling and color changes, which is laborious, time-consuming, and subjective [5].

The rapid advancement of optical sensors and imaging technologies has significantly impacted agriculture and brought about more automation [6]. Initially, imaging techniques used red-green-blue color systems for the identification of color change and defects in the food and agricultural products [7]. Since, multispectral fluorescence imaging has been used with maize, peas, soybean for measuring color change [8].

Hyperspectral Imaging (HSI) has also evolved over the years and is being explored as technique for nondestructive food analysis. HSI provides both spatial and spectral information about an object. HSI consists of many thousands of pixels in a two-dimensional array, with each pixel containing a spectrum corresponding to a specific region on the surface of the sample. These spectra vary according to unique material and chemical compositions. Interrogation of these spectra makes possible the development of mathematical models to estimate the chemical composition or functional class of a sample associated with each pixel. Results reported in several studies have indicated that hyperspectral imaging is able to predict a number of food components and quality parameters in a wide range of biological matrices [5]. Previous research has shown that HSI was used in a plethora of applications in agriculture and food industry to measure the textural changes including bruise, chilling injury, firmness [9,10,11], and biochemical detection such as moisture content, soluble solids content, acidity, and phenolics [12, 13,14], biosafety measurement in bacterial or fungal infection and fruit-fly infestation [15,16]. In addition, photosynthetic rate in mangroves was studied in relation to salinity stress using HSI technology [5]. More importantly, Hernandez et al. [17] reported that hyperspectral imaging can quantify the localization and total glucoraphanin in florets and stalks of broccoli. Moreover, the low instrument cost and fast-detecting properties of HSI have enabled the development of powerful diagnostic tools for detecting, classifying, and visualizing quality and safety attributes of fresh produce.

This study was conducted to evaluate the potential significance of hyperspectral imaging as a tool to determine the freshness of broccoli during postharvest storage. Through High Performance Liquid Chromatography (HPLC) quantitation, we showed that there is a linear correlation between the total glucosinolates concentration and post-harvest senescence in broccoli. Moreover, we performed Real-Time Polymerase Chain Reaction (PCR) for expression studies on 13 enzymes are involved in the biochemical pathway producing glucosinolates. We believe that combination of HSI and glucosinolates analysis can define the freshness signature in postharvest broccoli.

## Materials and methods

### Tissue collection and Preparation

Freshly grown broccoli (cultivar, Emerald Crown) florets were manually harvested from local farms in Hastings, Florida to avoid any mechanical damage. All the broccoli florets were selected with the same shape and size in this study. The florets were then stored in either a cold room (4–5°C, darkness) for the cold treatment, or in a plant growth chamber at 25°C (RT) with 16 hours of light and 8 hours of dark. Four biological replicate of broccoli florets were used for the experimental sampling for tissue collections. Tissue samples were collected from the broccoli florets every other day during a twelve-day period. The samples were wrapped in aluminum foil, immediately frozen in liquid nitrogen and then stored in −80°C for quantitative PCR and freeze-dried for HPLC.

### Hyperspectral imaging setup

Broccoli hyperspectral images were collected with a HinaLea 4200 hyperspectral camera and converted to reflectance spectra. The camera covers the wavelengths ranging from 400nm to 1000nm with a resolution at 4nm (based on full width at half maximum), resulting in 300 wavebands. Halogen lamps were used as illuminators in an imaging chamber. Within the chamber, each broccoli sample was placed on a black plate with matte black siding to absorb redundant light in order to minimize scattering. Hyperspectral measurements of three biological replicates were carried out at every time point and the experiment was repeated three times.

### Pre-processing Reflectance Spectra

Pre-processing of the measured spectra for noise and illumination variation was required prior to analysis. The measured reflectance spectra consistently contained higher levels of noise at the two ends of the wavelength range. In addition, since the illuminator used was a point light source and did not evenly cover the entire imaging surface. As a result, the center of the imaging plane was brighter than the outer edge. Fig. 1 (d) shows examples for the issues mentioned above. To mitigate these issues, first, a median filter of length 5 along the wavelength axis was applied to the reflectance spectra. Next, responses below 500 nm and above 900 nm were removed due to the high noise levels. Finally, in order to reduce the magnitude difference caused by point light source, we applied *l*_2_ normalization in which each spectral signature is normalized to have unit norm.

**Fig 1.**
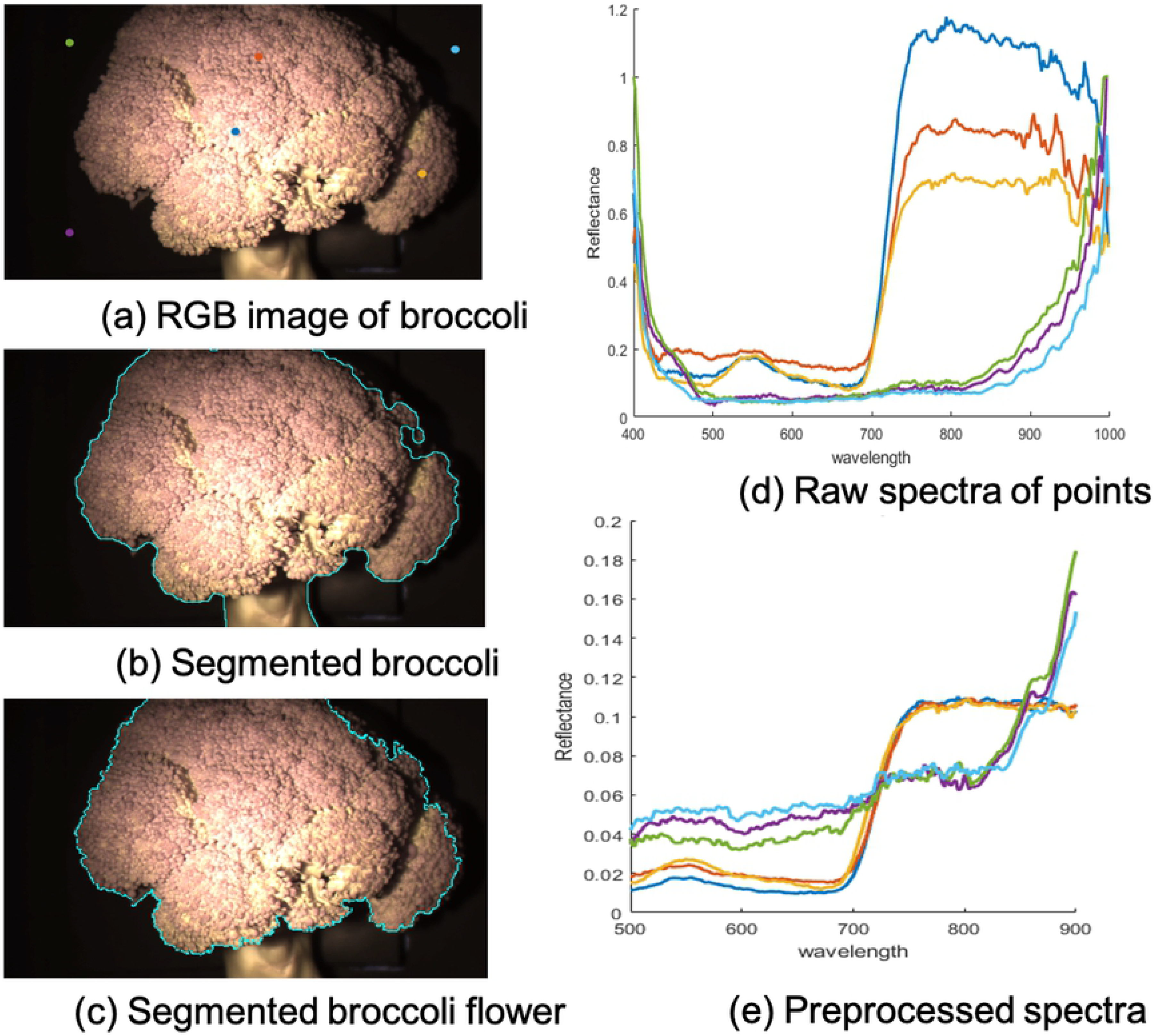
RGB image and spectra. (a) Broccoli sample placed on a black plate. (b) Segmented broccoli sample. (c) Segmented broccoli floret. (d) Reflectance measured by HinaLea 4200 hyperspectral camera. (e) Preprocessed spectra. The color of spectra in the (d)-(e) is corresponding to the points in (a).

### Segmenting Floret from Background

After pre-processing spectra, the regions of florets were segmented from the remainder of the hyperspectral cube. This segmentation was accomplished in two steps: (1) segmenting the broccoli sample from the black background; (2) segmenting the floret from the stalk. For the first stage, Fig. 1 (a) and (d)-(e) illustrates the spectral differences between the broccoli and the black background. Namely, the broccoli spectra have a bump around 550nm wavelength (visible bands of green) and a sharp increase around 700nm wavelength (Near infrared/Red edge whereas the spectra for the black background are nearly flat up to 800nm and then increases rapidly. Given these significant spectral differences, the k-means clustering algorithms was applied to cluster the pixel spectra into two groups, broccoli and background. After clustering, a morphological image closing operation was applied to connect any disconnected points. An example of the segmented results is shown in Fig. 1 (b).

Since the glucosinolates concentration was measured on broccoli spores, we hypothesized that focusing on the spectra of broccoli flower will generate stronger correlation than analyzing the spectra of entire broccoli. In order to segment the floret from the stalk, we applied the GraphCut algorithm [18] of the image segmentation toolbox in Matlab 2019b [19]. The algorithm was seeded by the user providing a marking that denotes the broccoli flower and the background including broccoli pedicle. The segmentation took around 5 to 10 seconds for each image. The segmented results are shown in Fig. 1 (c).

### Unmixing and correlation with glucosinolates information

The hyperspectral image collects a high-dimensional image cube that describes each pixel as the radiance or the reflectance at a range of wavelengths across the electromagnetic spectrum. The spectrum of a pixel is usually determined by the material of the object surface. With the assumption that the measured spectrum is consists of a set of constituent spectra, also known as endmembers, spectral unmixing is defined as decomposing the mixed spectrum into a collection of endmembers and their corresponding proportions, also known as abundances [20]. A well-known spectral model (and the most commonly applied to perform hyperspectral unmixing) is the linear mixing model (LMM) which represents each measured spectrum as a convex combination of endmembers as shown in Equation 1 [20],

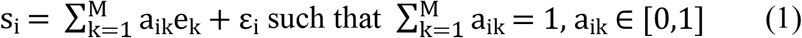

where s_i_ is the spectrum of pixel i, ε_i_ is the noise vector, M is the number of endmembers, e_k_ is k^th^ endmember, and a_ik_ is the corresponding abundance value. The objective of each unmixing broccoli sample is to find a set of endmembers and abundances that can represent the freshness of broccoli. Thus, in this work, we attempt to estimate endmembers that represent the range of “freshness” levels in the samples. Then the associated abundances for the endmembers corresponding to “fresh” samples can use viewed as a freshness indicator for the sample. The endmembers and abundances were estimated using two approaches averaging (as described below in Section A) or the application of the SPICE algorithm (as described in Section B).

#### A. Unmixing with averaging spectra as endmembers

In the averaging approach, the endmembers were extracted by averaging the broccoli spectrum from two extreme storage conditions. Specifically, ***e***_1_ is the average from broccoli on day 1, representing the best fresh level. ***e***_2_ is the average from broccoli kept under room temperature for 12 days, representing the least fresh level. The greater *a*_*i*1_ indicates more fresh level of pixel *i* in the broccoli sample.

The abundance values were then estimated by optimizing the fully constrained least squares of each pixel with the above two endmembers as shown in Equation 2 [22].

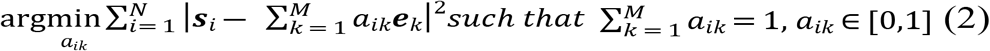

where *M* = 2 in this case.

#### B. Unmixing with SPICE algorithm

Instead of extracting average spectra as endmembers, the sparsity promoting iterated constrained endmember (SPICE) algorithm benefits from simultaneous estimating the shape and number of endmembers as well as the abundances [21]. The algorithm is initialized with a large number of endmembers and iteratively updates the estimated endmembers and abundances by optimizing Equation 3 using an alternating optimization approach,

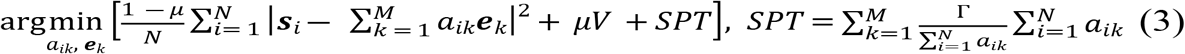

where *V* is the sum of variances among estimated endmembers, *μ* is the regularization parameter to balance the reconstruction error and variance, Γ is a constant that decide the proportion values are driven to zero, and *a*′_*ik*_ is the abundance value for the *i*th pixel in the *k*th endmember from the previous iteration. After each iteration, endmembers which are not being used to represent the data are pruned from the overall endmember set^1^.

#### C. Correlating abundance with glucosinolates concentration level

The estimated abundance vectors 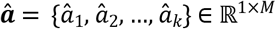 for each broccoli sample (where the value 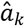 is the average abundance of all pixels over the region of interest as 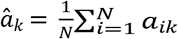, where *N* denotes the number of pixels) were used to attempt to predict measured glucosinolates concentration values. Specifically, a multi-variable linear regression (MLR) model as Equation 4 was fit using least squares estimation approaches to predict the glucosinolates concentration value,

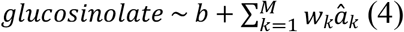

where *b* is the bias, *w*_*k*_ is the coefficient for 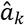, *M* is the number of estimated endmembers.

### Extraction of total glucosinolates for HPLC quantification

HPLC-UV analysis of total glucosinolates was extracted according to previously defined methods with some modifications [22]. Raw materials from previously harvested broccoli florets were stored at −80°C and further samples were taken out to be dispersed in liquid nitrogen. 100 mg of samples was weighed and crushed to fine powder using mortar and pestle. Subsequently, crushed tissue powder was dissolved in 1 ml of 50 % methanol contained in the 1.5 ml eppendorf tube. The tubes were further kept at 65°C for 1 hour in the water heating bath. Samples were then centrifuged at 15000 g for 10 minute. The supernatant was filtered through a 0.22um hydrophilic PTFE syringe filter (Sigma Aldrich, USA). HPLC analysis was done with the flow rate of 0.4 ml/min at a column temperature of 40°C with a wavelength of 227 nm. The solvent used were water and 100 % acetonitrile. The individual glucosinolates were estimated by their HPLC peak area with reference to a desulfo-sinigrin method [22]. Total peak area was calculated from broccoli florets from day 1, 3, 5, 7, 9, 11 when stored at growth chamber (25°C) and cold temperature (4°C). Four biological replicate were used for each time point during the study.

### RNA extraction and gene expression studies using Quantitative real-time PCR

Florets tissue samples from day 1, 3, 5 were chosen for glucosinolates pathway expression analysis. Total RNA was isolated from broccoli floret tissue stored in liquid nitrogen using TRIZOL (Ambion, life Technologies) method and DNase treatment (Turbo DNA free, Thermo-fisher). First strand cDNA synthesis with 1μg of total RNA was performed using reverse transcription kit (Applied Biosystems). For quantitative real-time RT-PCR, primers were designed using Primer Quest, Integrated DNA Technologies (IDT) software. The primers of glucosinolates pathway genes were listed in Table 1. Real time PCR reaction was performed in Applied Biosystems qPCR machine (Thermofisher). Total reaction was 10 μl for each gene in triplicates with thermocycler conditions as: 95°C for 10 min, 45 cycles for 95°C for 30 sec, 60°C for 30 sec. Relative gene expression was calculated by ΔCT method. Actin 2 was used as an internal control. This experiment was repeated twice.

**Table 1 :**
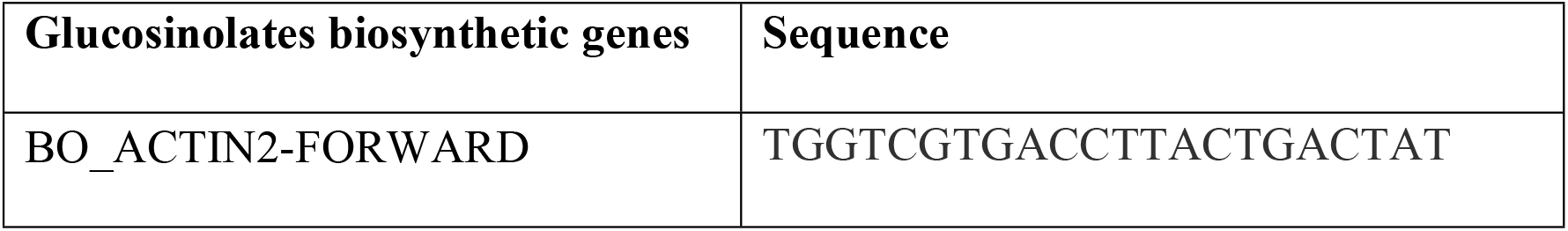
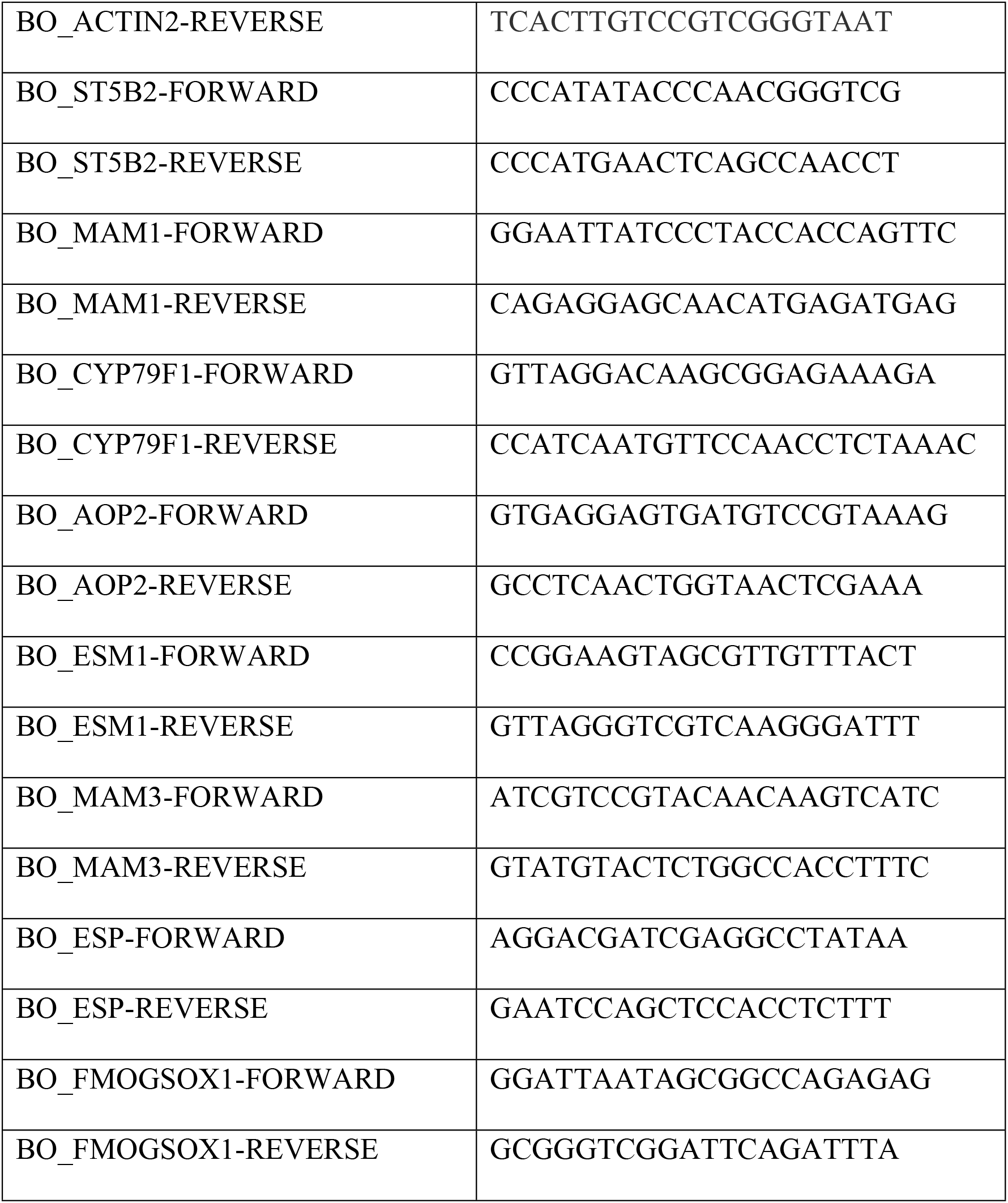
List of primers for quantitative PCR performed for glucosinolates pathway in Broccoli.

## Results

### Use HSI as tool to detect postharvest senescence in broccoli

First, spectra were downsampled to 5,000 per segmented broccoli sample via k-means clustering to accelerate computing. Downsampling was applied to the two segmentation scenarios being considered: (1) entire broccoli, and (2) broccoli florets, respectively. Next, the entire dataset was split into training, validation and testing sets. To be more specific, 48 samples under 12 conditions (2 storage conditions over 6 time stages) were randomly divided into 4 folds, one for testing, and the other 3 for training and validation. Each fold contains 12 samples, 1 replication from each condition. The training and validation dataset were shuffled in every repetition.

In the first step, two endmembers were calculated by averaging spectra from the most and least fresh replication in the training folds. The glucosinolates concentration and derived abundance feature for all training replications were applied to fit the MLR model. The training process was repeated for 10 times over 3 folds. In each repetition, we tested the trained model on the testing fold, calculated the mean and standard deviation of the testing and training prediction error in Table 2. In addition, a model was selected according to the root means square error (RMSE) and R-squared value from training and validation folds and was applied to the testing fold to generate the result shown in Fig. 2.

**Table 2.**
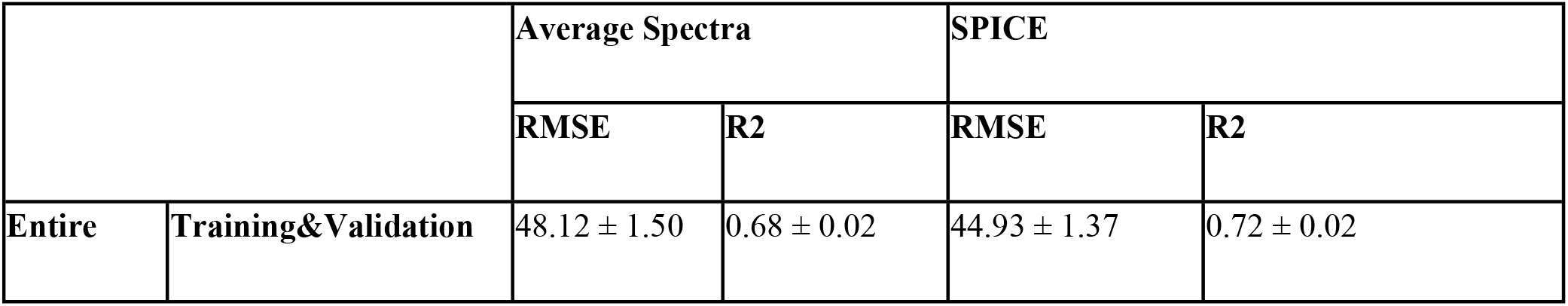
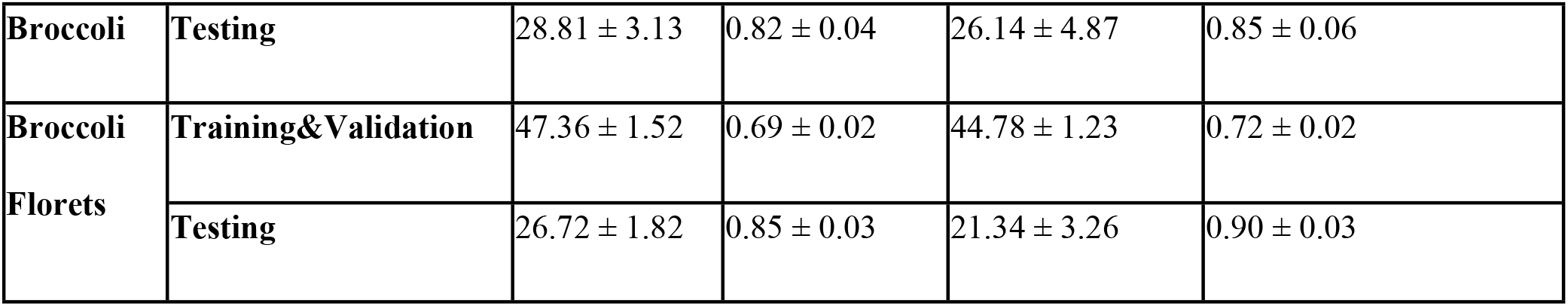
Comparison of glucosinolates prediction error from endmember abundances.

**Fig 2.**
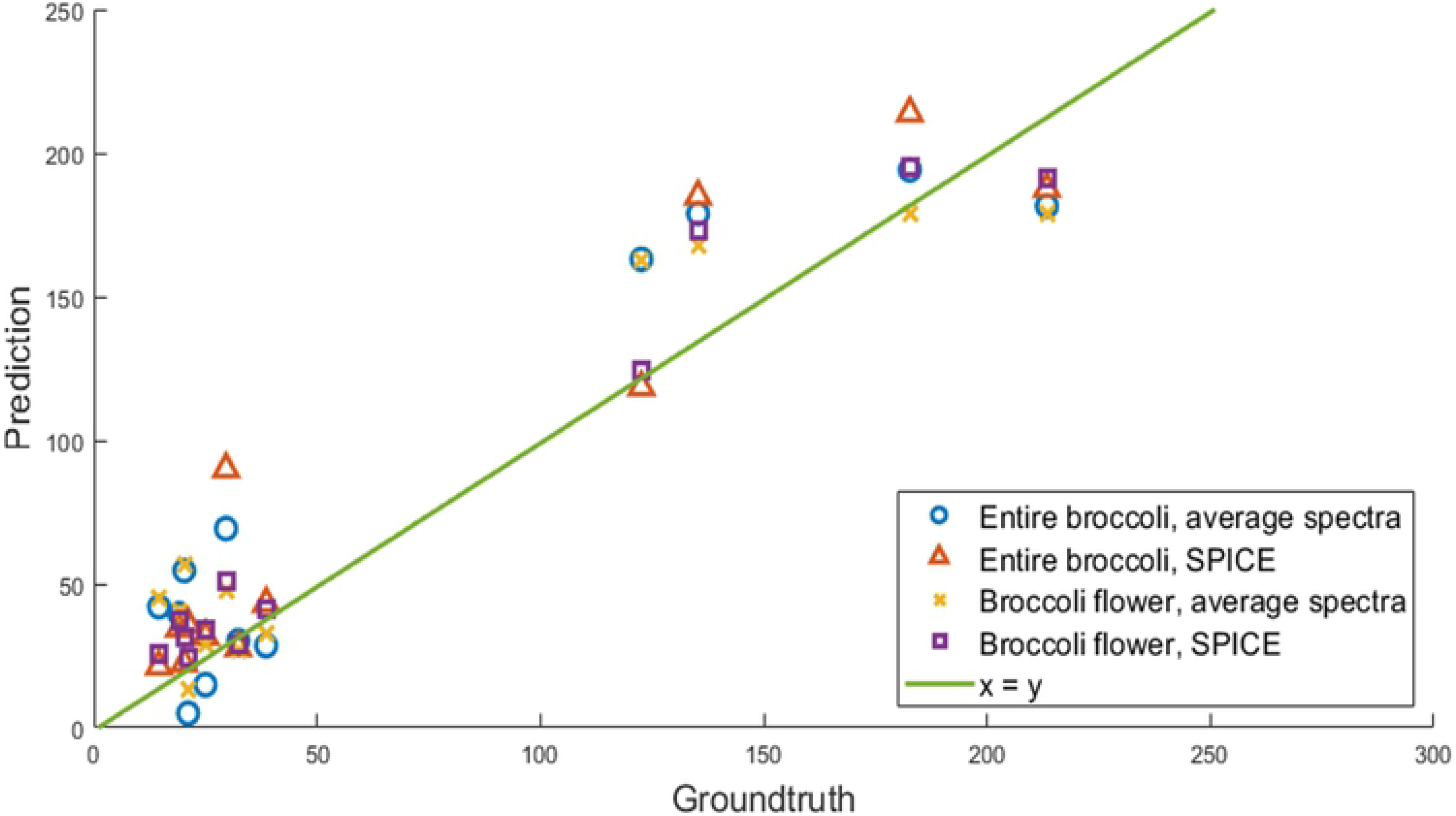
Predicted glucosinolates on testing fold. x-axis indicates the real glucosinolates concentration, y-axis indicates the prediction. Markers in various shape and color denotes prediction with different methods. Markers that closer to the “x = y” line indicates more accurate prediction.

Next, endmembers were estimated from training folds via SPICE. Since the estimated endmembers highly depends on the parameter Γ, a various range of Γ, starting from 10 to 150 with stepsize 10, were explored. We conducted 10 repetitions over 3 folds for 15 Γ values. Similar to the first stage, with the estimated endmembers, the abundance feature can be derived from the training folds to fit the MLR model. Fig. 3 (a-b), (f-g) shows greater prediction error on validation folds with a smaller Γ, which indicates over-fitting of the approach. Namely, Since Γ determines the proportion of estimated endmembers to be eliminated, a smaller Γ results in a greater number of endmembers and more parameters that need to be estimated (and provide opportunity for overfitting). Fig. 3 (c-d), (h-i) illustrate the tendency of over-fitting with increasing number of endmembers *M*. In addition, according to Fig. 3 (e), (j), *M* = 3 was determined for both segmentation, since it is the most number over 450 replications. We tested the trained MLR models with estimated endmembers on the test fold, generating the prediction errors shown in Table 2. A model was selected via the same criterion from repetitions and was applied to the same test fold as the first stage. The prediction results are shown in Fig. 2.

**Fig 3.**
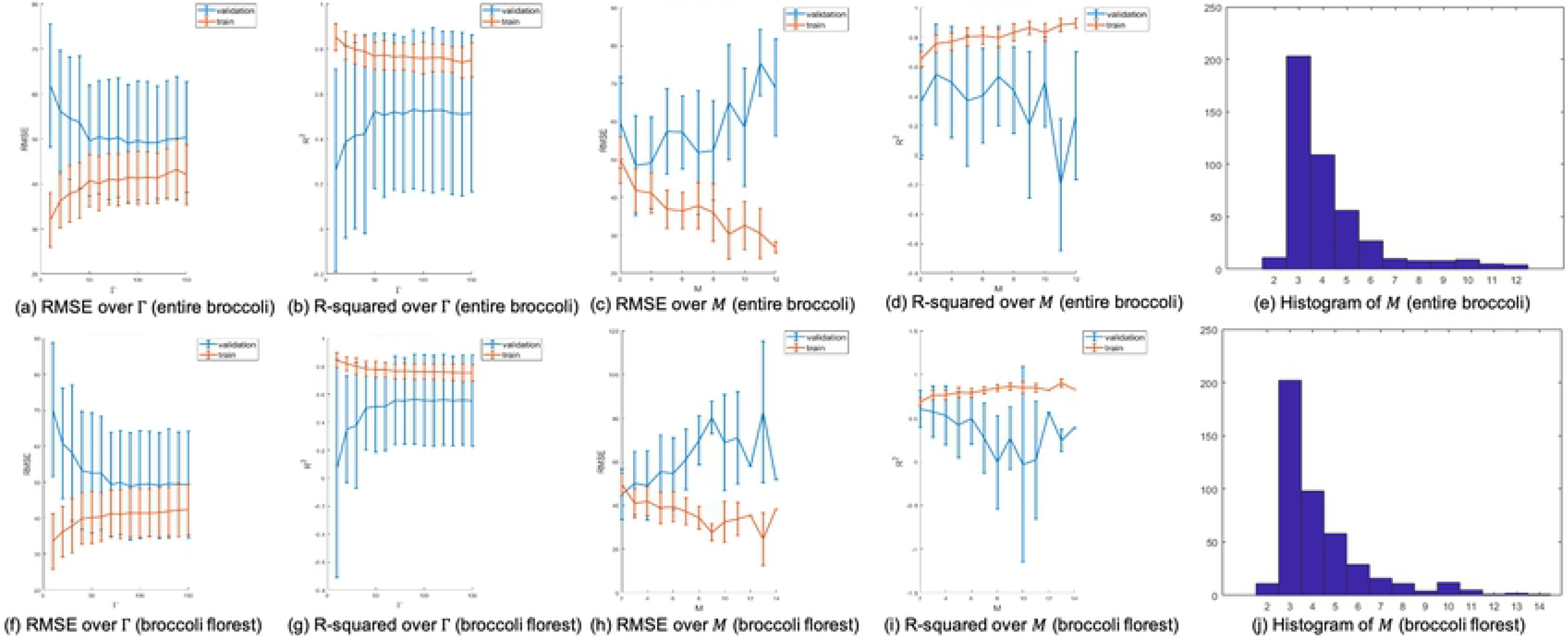
Exploration of SPICE parameters. (a-e) The training and validation error across various parameter setting for broccoli florets. (f-j) The training and validation error across various parameter setting for entire broccoli replicate.

The visualization of unmixing result is shown in Fig. 4, where (a-b) plot the estimated endmembers with average and SPICE on broccoli florets. Correspondingly, (c-d) visualize the estimated abundance map for testing samples in day 1, day 5, and day 12, and (e-f) plot the histogram of abundance value. The distribution of abundance and the number of pixels associated with each endmember is informative of freshness over days. In addition, the abundance map for the 3rd endmember in SPICE shows a relatively high concentration in day 5. It would be worth to explore whether the 3rd endmember can reveal a transition status during the progress of decay.

**Fig 4.**
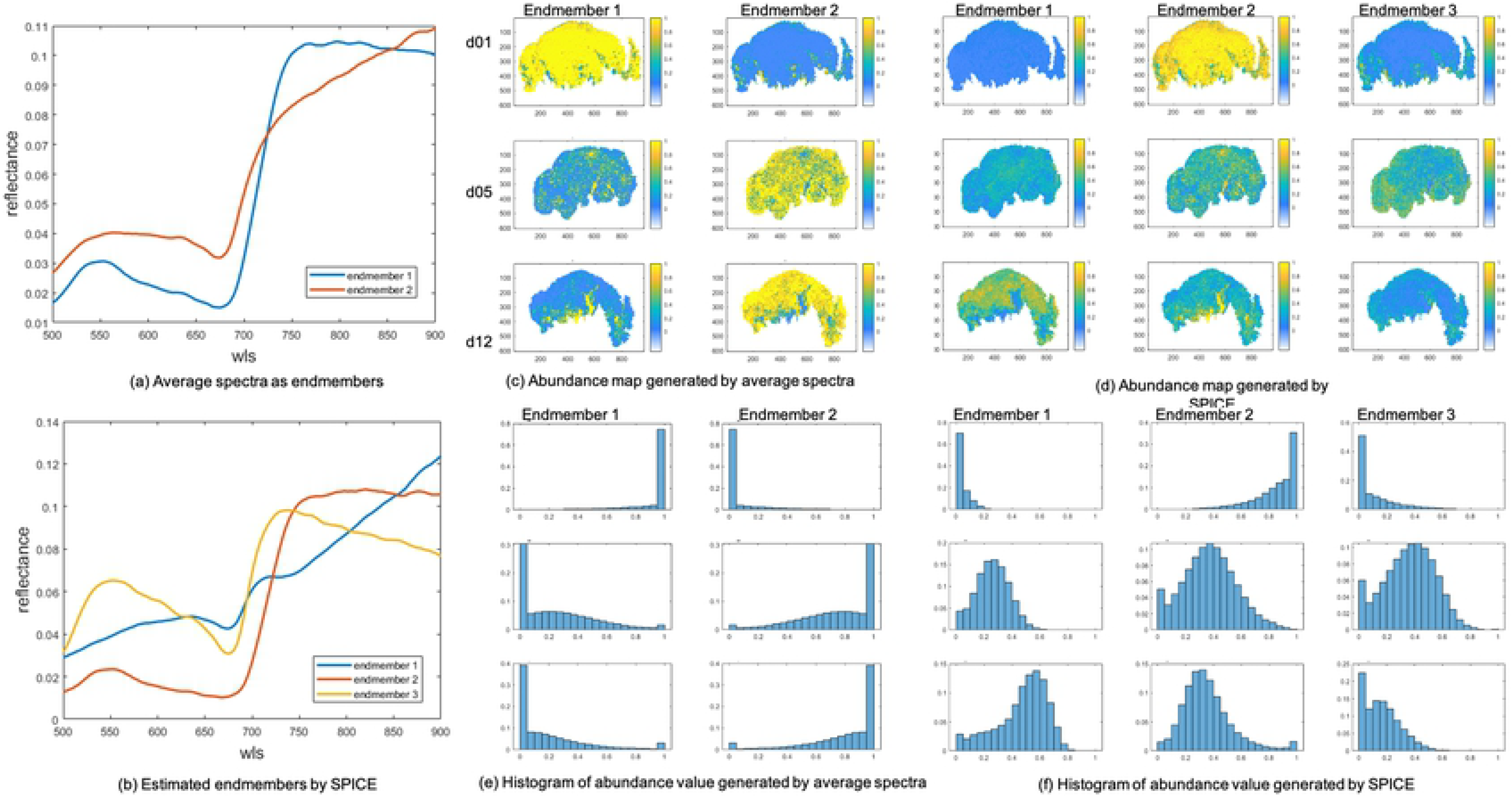
Estimated endmembers and unmixing results of testing samples in day 1, 5 and 12. (a-b) Estimated endmembers with average and SPICE on broccoli florets. (c-d) Visualization of estimated abundance map for testing samples in day 1, day 5, and day 12. (e-f) Histogram of abundance value.

### Indo-glucosinolate peaks increased during the postharvest senescence

To access the possibility that the glucosinolates can be applied as senescence indicator, we performed HPLC analysis to measure the total glucosinolates concentration during the postharvest stored broccoli. In HPLC analysis, we monitored total peak area for indole-glucosinolates, the most widely distributed glucosinolates, at six time points during a 12-day of period for the broccoli florets stored at room temperature (25°C) and cold temperature (4°C). Four biological replicates were analyzed at each timepoint. We found that there is a linear increase in indo-glucosinlates concentration throughout a 12-day period in broccoli when stored at room temperature (Fig 5). However, this changes were not observed during the cold storage treatment in broccoli. Thus, there is a strong correlation observed between the indo-glucosinolates levels and progression of postharvest senescence in broccoli. This data suggested that indo-glucosinlates can potentially serve as an ‘freshness indicator’ to define a freshness signature.

**Fig 5 .**
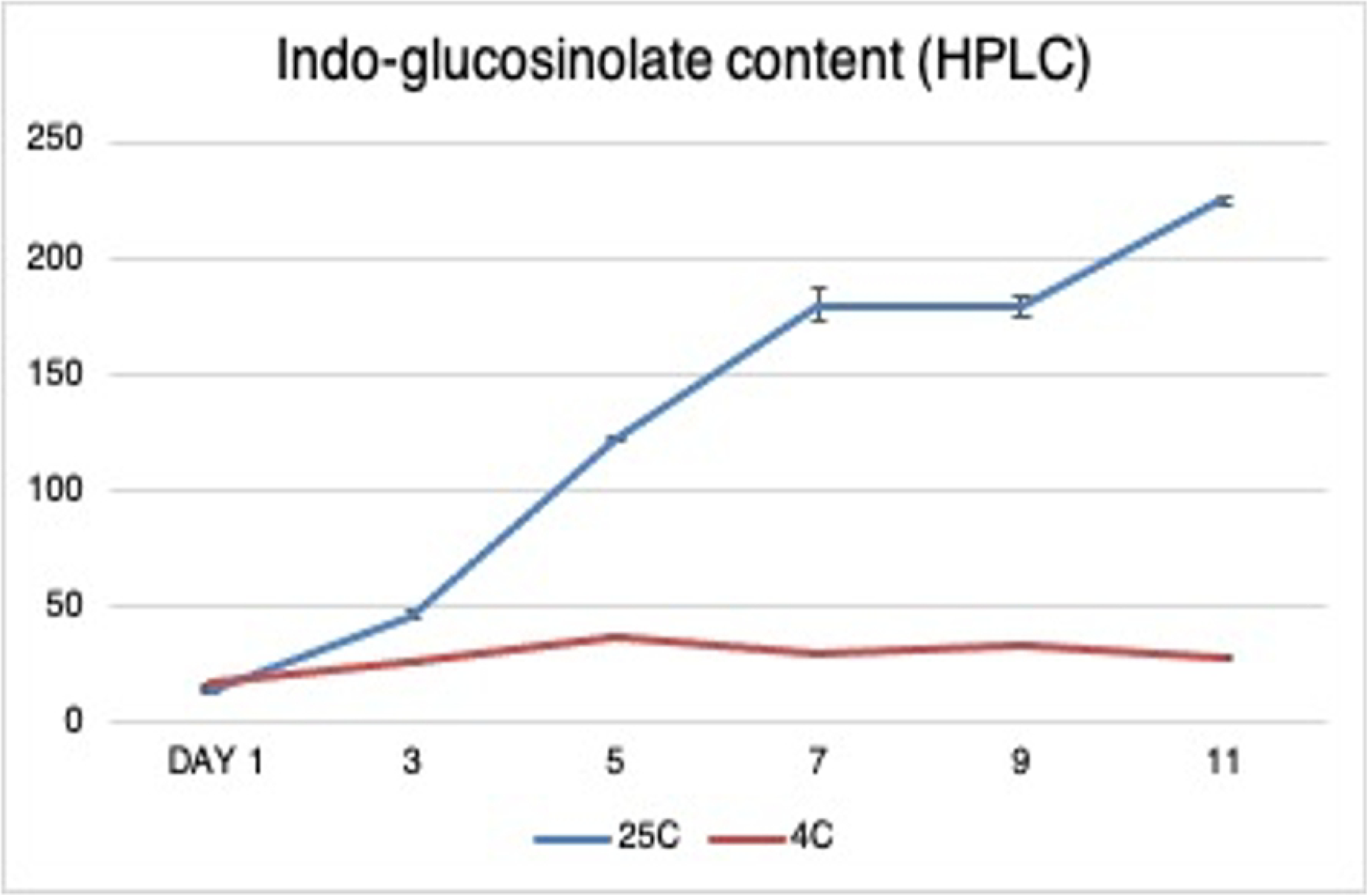
Quantification of the indo-glucosinolates peak area by HPLC. Graph displayed the changes in the indo-glucosinolates level in the broccoli florets on day 1, 3, 5, 7, 9, 11. Data represented means ± SE bars (n=4 for each day).

### Glucosinolates transcriptional levels were increased during the postharvest senescence

To further validate this correlation and examine how the glucosinolates biosynthetic pathway is affected during the postharvest senescence process, we carried out quantitative gene expression of key genes in the glucosinolates biosynthetic pathway and observed that the expression of all genes involved in glucosinolates biosynthetic pathway were up-regulated when stored at room temperature. Mostly, all the key candidates’ genes showed significant increase in transcript expression on day 5 when the broccoli florets were stored at room temperature however, the increase in glucosinolates profile during the postharvest day 5 was significant but less drastic than the senescent conditions (Fig 6). This data showed that glucosinolates levels increased rapidly in room temperature when stored at the room temperature. In case of pathway intermediate methylthioalkylmalate synthases (MAM1 and MAM3), there was 4.3 fold and 11 fold significant increase in transcript levels from day 1 to day 3 and day 3 to day 5, respectively at room temperature. However, in cold conditions, MAM1 levels were undetectable on day 3 but increased on day 5 by 5.3 fold (Fig 6). This implied that MAM1 levels were increased in higher proportions under room temperature condition. Similar patterns were observed for MAM3, *epithiospecifier modifier 1* (*ESM1*), α-ketoglutarate-dependent dioxygenase (*AOP2*), epithiospecifier protein (*ESP2*), CYP79ST5B2 as their transcripts were significantly higher from day 0 to day 5. However, under cold conditions, the increase in flavin-monooxygenases (FMOGSOX2) transcript levels from day 0 to day 5 was not significant. This observation provided evidence that the gene expression changes in glucosinolates pathway were associated with postharvest storage conditions. Transcript levels for all genes were significantly higher at 25°C indicating that the cold temperature was inhibited indo-glucosinolate production in postharvest broccoli. Our results suggested that there is correlation between senescence and indo-glucosinlates concentration in postharvest broccoli.

**Fig 6.**
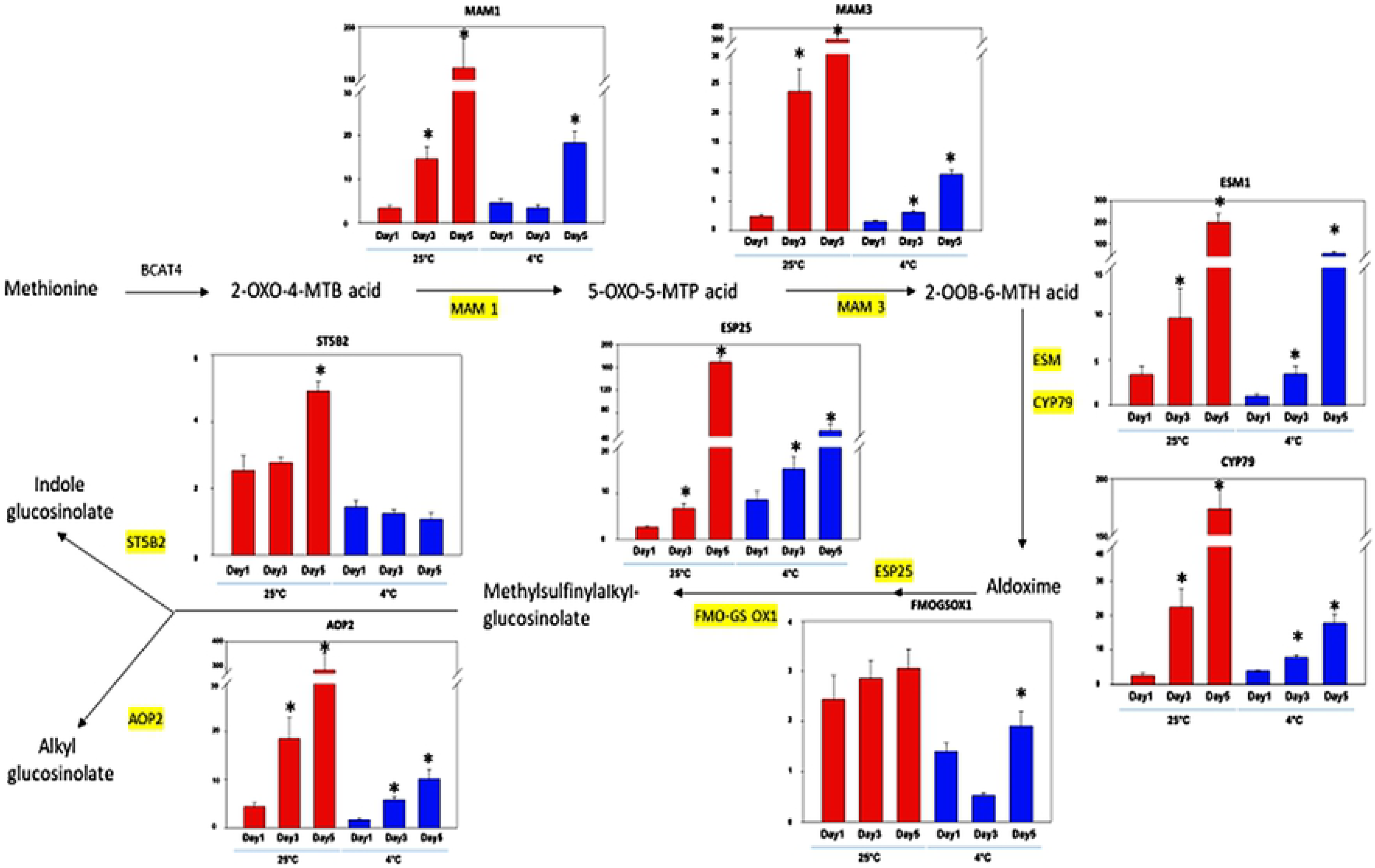
A flowsheet of Indo-glucosinolates biosynthetic pathway displaying the transcript levels on day 1, 3, 5 for key enzymatic intermediates catalyzing the biosynthetic pathway.

All the key genes in glucosinolates biosynthetic pathway are highlighted in yellow. Red bars in the graph at each step shows the transcript level of specific candidate genes at 25°C on day 1, 3, 5 whereas blue bars show the expression level of the genes at 4°C on day 1, 3, 5. Expression of each gene was normalized using actin as an internal control. Data represents means ± SE bars (n=3). Asterisks (*) indicate statistically significant differences from day 1 (control) to day 3, day 5 (storage temperature conditions) (*p* < 0.05).

### Outliers

The smaller prediction error for testing fold compared with training and validation folds in Table 2 can be explained by the observed outliers S1 Fig. showed the prediction performance of the training fold with SPICE on the broccoli florets. The marker size and color are related to their prediction error. Apparently, the three circled outliers generated greater prediction error compared with others. The S1 Table listed the glucosinolates concentration of 4 replications kept in room temperature over 6 time points. Three bold numbers are corresponding to the circled outliers in S1 Fig. In rep 1, the glucosinolates concentration was increasing along days, while in rep 2-4, the bold number showed “abnormal” performance. An additional experiment was conducted, where the outliers were moved to the testing fold. S2 Fig. and S2 Table showed the prediction performance and error on the additional testing fold.

## Discussion

The above experiments were conducted on both the entire broccoli sample and the segmented broccoli florets. Fig. 4 and Table 2 compared the prediction performance and error. Overall, results depicted that the estimated abundances can indicate change in the glucosinolates concentration values. The prediction error can be explained by the fact that the measurement of hyperspectral data and glucosinolates concentration were conducted across different sample scales. Namely, the abundance values are derived from imaging across the entire surface of one side of a broccoli sample, whereas the glucosinolates value is measured using only one small component of the broccoli tissue. The RMSE values show that unmixing with broccoli florets only has slightly less error than when using the entire broccoli sample. In addition, SPICE outperforms the simple averaging. However, when considering the computing and operation complexity, averaging spectra is a more straightforward approach to estimate endmembers as compared with SPICE and does not require parameter selection.

In summary, hyperspectral imaging holds promising strength in demonstrating state of art performance in the area of crop sciences through the modulation of imaging with spectroscopy. As shown in this effort, HSI has the potential to provide quantitative parameters in understanding postharvest senescence.

## Acknowledgements

We thank Dr. Diane Rowland for providing the HinaLea 4200 hyperspectral camera and converted to reflectance spectra for supporting in sample imaging. We thank Dr. Ru Dai and Dr. Jeongim Kim for the HPLC analysis and troubleshooting. This work was supported by UF Seed Fund (#P0175583 to A.Z., T.L.).

## Supporting information

**S1 Fig. Predicted glucosinolates on training folds.** x-axis indicates the real glucosinolates concentration, y-axis indicates the prediction. Markers that closer to the “x = y” line indicates more accurate prediction. The marker size and color is corresponding to the prediction error, the bigger and brighter markers indicate greater error.

**S2 Fig. Predicted glucosinolates on additional testing fold**. x-axis indicates the real glucosinolates concentration, y-axis indicates the prediction. Markers in various shape and color denotes prediction with different methods. Markers that closer to the “x = y” line indicates more accurate prediction.

**S1 Table.** The glucosinolates concentration under 25°C over days.

|  | **Rep1** | **Rep2** | **Rep3** | **Rep4** |
| --- | --- | --- | --- | --- |
| **Day1** | 11.9544 | 11.9405 | 14.2901 | 21.099 |
| **Day3** | 53.858 | 31.6877 | 75.4149 | 25.0054 |
| **Day5** | 129.0532 | 113.1505 | 122.6086 | 122.941 |
| **Day8** | 135.2934 | 151.8804 | **348.884** | **82.8482** |
| **Day10** | 182.7224 | **80.5022** | 192.528 | 261.347 |
| **Day12** | 200.8352 | 269.094 | 215.7358 | 213.3986 |

**S2 Table.**
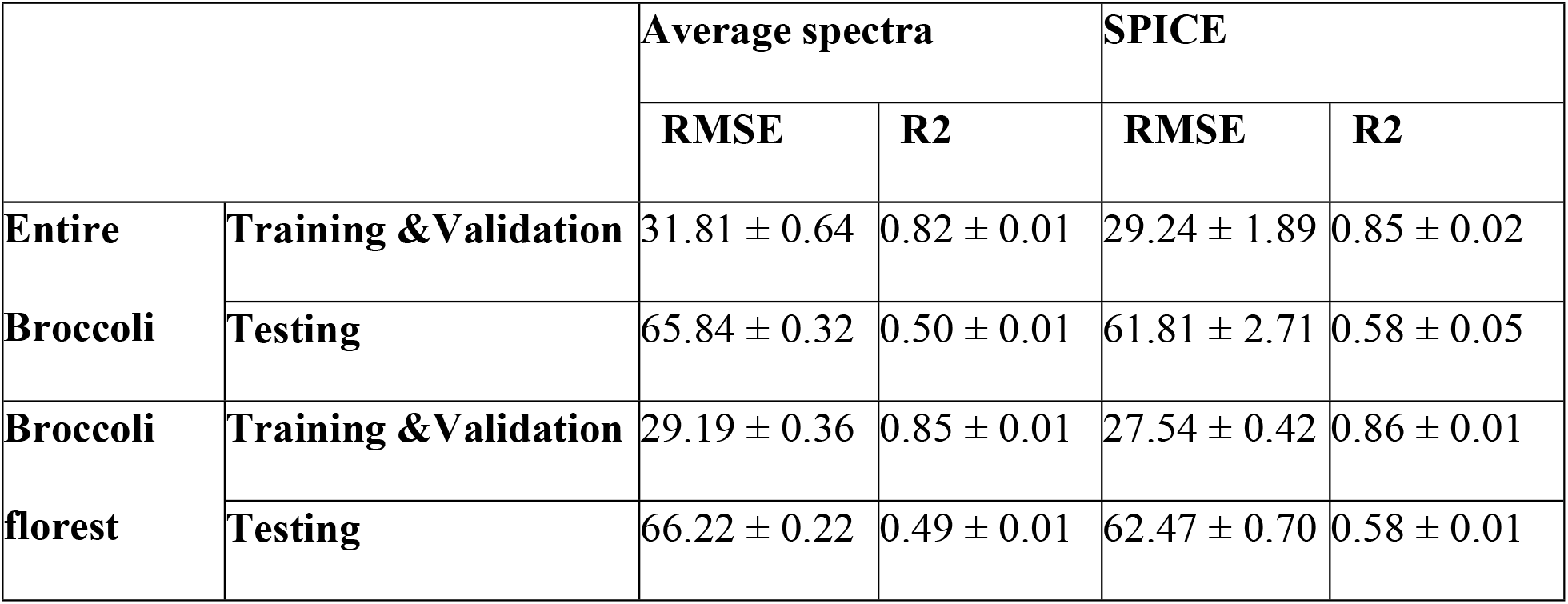
Comparison of prediction error on additional testing fold.

## Footnotes

1 The Matlab and Python implementation for SPICE can be found here: github.com/GatorSense

